# Refgenie: a reference genome resource manager

**DOI:** 10.1101/698704

**Authors:** Michal Stolarczyk, Vincent P. Reuter, Neal E. Magee, Nathan C. Sheffield

**Affiliations:** Center for Public Health Genomics, University of Virginia; Department of Public Health Sciences, University of Virginia; Department of Biomedical Engineering, University of Virginia; Department of Biochemistry and Molecular Genetics, University of Virginia; Research Computing, University of Virginia

## Abstract

Reference genome assemblies are essential for high-throughput sequencing analysis projects. Typically, genome assemblies are stored on disk alongside related resources; for example, many sequence aligners require the assembly to be *indexed*. The resulting indexes are broadly applicable for downstream analysis, so it makes sense to share them. However, there is no simple tool to do this. To this end, we introduce refgenie, a reference genome assembly asset manager. Refgenie makes it easier to organize, retrieve, and share genome analysis resources. In addition to genome indexes, refgenie can manage any files related to reference genomes, including sequences and annotation files. Refgenie includes a command-line interface and a server application that provides a RESTful API, so it is useful for both tool development and analysis.

**Availability:** https://refgenie.databio.org

## Background

Enormous effort goes into assembling and curating reference genomes^1–5^. These reference assemblies provide a common representation for comparing results and they form the basis for a wide range of downstream tools for sequence alignment and annotation. Many tools that rely on reference assemblies will produce independent resources that accompany an assembly. For instance, many aligners must *hash* the genome, creating *indexes* that are used to improve alignment performance^6–9^.

Analytical pipelines typically rely on these aligners and their indexes for the initial steps of a data analysis. These assembly resources are typically shared among many pipelines, so it’s common for a research group to organize them in a central folder to prevent duplication. In addition to saving disk space, centralization simplifies sharing software that uses a reference assembly because software can be written around a standard folder structure. However, this does not solve the problem of sharing genomic resources *between* research groups. Because each group may use a different strategy to identify shared genome resources, sharing tools across groups may require modifying them.

One solution to this problem is to have a web-accessible server where standard, organized reference assemblies are available for download. Indeed, this is exactly the goal of Illumina’s *iGenomes* project, which provides “a collection of reference sequences and annotation files for commonly analyzed organisms^10^.” The iGenomes project has become a popular source of genome assets and has greatly simplified sharing analysis tools among research environments. However, this approach suffers from some fundamental drawbacks and leaves several challenges unsolved. First, the individual assets can only be downloaded in bulk, but what if a particular use case requires only a small subset of resources in a package? More important, building the resources is not scripted, so if the repository excludes a reference or resource of interest, there is no programmatic way to fill the gap. In these scenarios, users must manually build and organize genome assets individually, forfeiting the strength of standardization among groups.

To improve the ability to share interoperable reference genome assets, we have developed *refgenie*, which enables a more modular, customizable, and user-controlled approach to managing reference assembly resources. Like iGenomes, refgenie standardizes reference genome asset organization so software can be built around that organization. But unlike iGenomes, refgenie also automates the *building* of genome assets, so that an identical representation can be produced for any genome assembly. Furthermore, refgenie allows programmatic access to individual resources both remote and local, making it suitable for the next generation of self-contained pipelines.

Refgenie can organize any files that can be assigned to a particular reference genome assembly, which could include not only genome indexes, but other resource types like genome sequences and annotations^11–13^.

Refgenie manages genome-related resources flexibly. It can handle any asset type, from annotations to indexes. It provides individual, pre-built asset downloads from a server and allows scripted building for custom inputs. Refgenie thus solves a major hurdle in biological data analysis.

## Results and discussion

Refgenie is the first full-service *reference genome asset manager*. Refgenie provides two ways to obtain genome assets: *pull*, and *build* (Fig.1A). For common assets, *pulling* a pre-built version obviates the need to install and run specialized software to build a particular asset. It also makes it easier to satisfy prerequisites programmatically for pipelines and other software. However, remote-hosted assets are only practical for common genomes and assets, so for uncommon assets or on unconnected computers, users may instead *build* assets, which creates the same standard output for custom genomes. By providing both *build* and *pull*, refgenie facilitates asset organization both within and between research groups, increasing interoperability of tools that rely on genome resources.

**Fig. 1:**
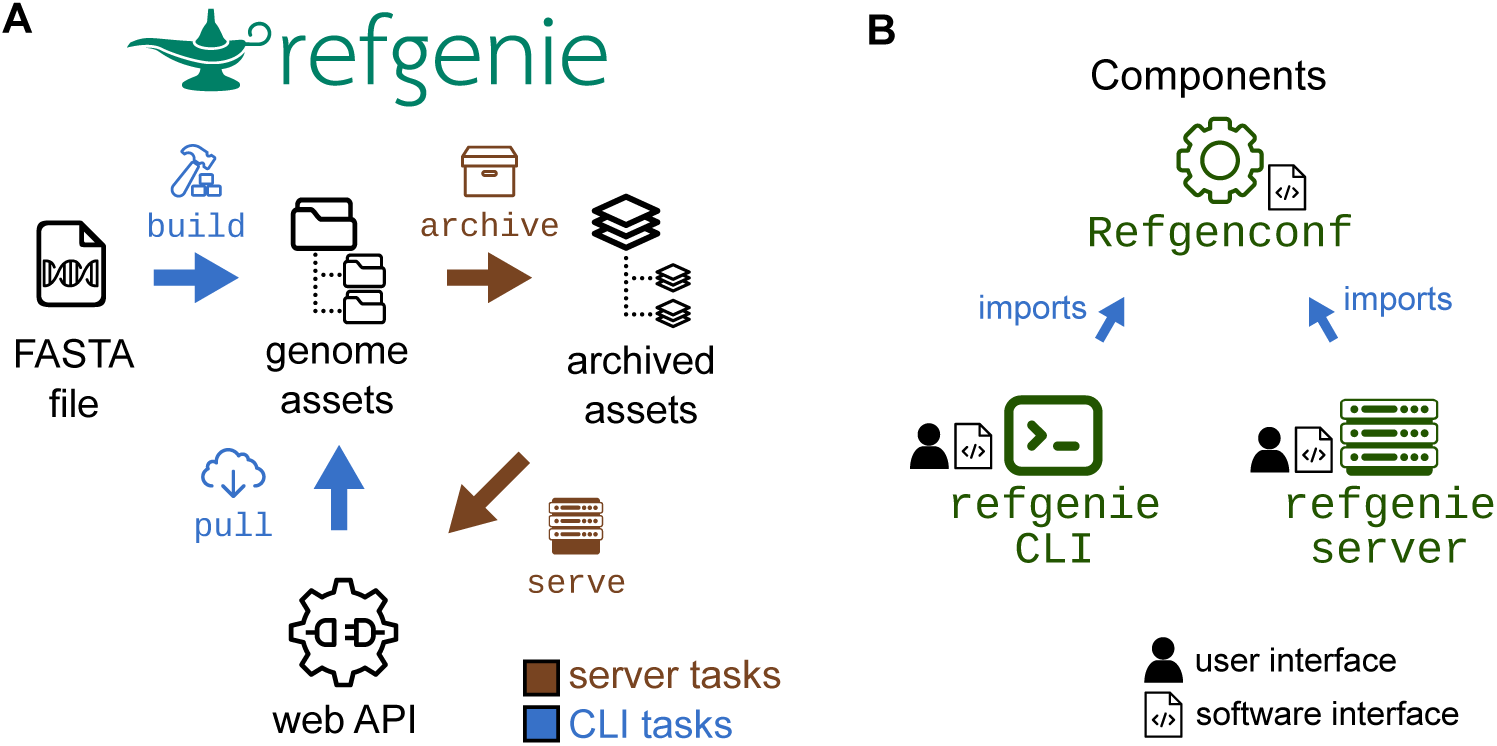
Refgenie concept and software organization. A: Refgenie provides the ability to either build or pull assets. B: Refgenie is tripartite, made up of a conf utility, a command-line interface (CLI), and a server package. The configuration package is intended for programmatic use, and is used by the CLI and server packages. Users and software use refgenie via the CLI or server (web API).

The `refgenie` software suite consists of three components: 1) a command-line interface (CLI), 2) a server, and 3) a configuration package that supports them both (Fig.1B). Each of these relies on a local YAML file called the *genome configuration file* (Fig. 2), which refgenie uses to keep track of metadata, such as local file paths.

**Fig.2:**
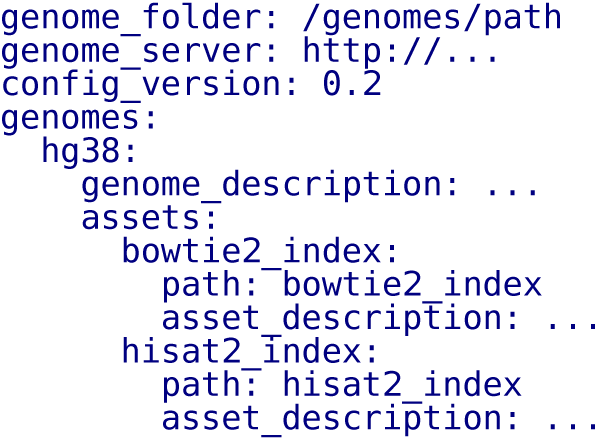
Genome config file. Refgenie reads and writes a genome configuration file in YAML format to keep track of available local assets.

### Genome configuration and asset organization

Refgenie organizes genome assembly resources into *assets*, each of which represents one or more files. You can think of a genome asset as a folder of related files tied to a particular genome assembly. For example, an asset could be an index for a particular tool, or a group of annotation files. Refgenie organizes such assets by genome in the configuration file, which is both computer-readable and human-readable. In practice, users will not need to interact with this file at all, as refgenie will handle both reading and writing the file. However, users may edit the file if they need a more complicated structure (such as storing assets on different file systems, or adding assets manually). Together with the refgenie software, this simple file makes the concept of reference genome assets completely portable. Full documentation for the configuration file format can be found at refgenie.databio.org.

### Refgenconf configuration package

The configuration package, `refgenconf`, simply provides functions and data types to read and write items listed in the genome configuration file.

Under the hood, the refgenie CLI itself uses `refgenconf` to interact with the genome configuration and assets on disk. The server software also relies on it to read, archive, and serve assets. The `refgenconf` package also provides the starting point for any third-party python developers by providing a fully functional python application programming interface (API) for interacting with refgenie assets. For example, we use `refgenconf` in python pipelines we develop to make them aware of the genome assets available in a given computing environment. Using this approach, a pipeline need only be provided with an assembly key, like ‘hg38’, and it can use `refgenconf` to locate the correct path to any genome-related asset necessary for the pipeline. This simplifies the process of configuring pipelines and allows `refgenie` to be used both by humans and computers.

### Refgenie command-line interface

The workhorse of refgenie is the command-line interface (CLI); it is how users will typically interact with genome assets. Its implementation as a command-line tool not only makes it useful for general purpose exploration and access, but also allows it to be integrated into existing workflows that require access to genome assets from the shell. The CLI can be installed with `pip install refgenie` and invoked by calling `refgenie`. The refgenie CLI provides 5 functions for interacting with local genome assets:

- `refgenie init` initializes an empty genome configuration file
- `refgenie list` summarizes the genome configuration file, listing local genomes and assets
- `refgenie seek` provides the file path to a given asset
- `refgenie add` adds an already-built local asset
- `refgenie build` builds a new asse

The `init, list, seek`, and `add` functions follow directly from the configuration file format. They essentially allow a user to easily explore and access file paths to available assets. The `build` function allows a user to *build* assets for any FASTA file, which is a more flexible system than alternative approaches that provide only downloadable assets. Refgenie has built-in capability to build a selection of different common genome assets (Fig.3). In addition to functions on local assets, the refgenie CLI also contains additional commands that can interact with remote assets: *pull* and *listr*:

**Fig.3:**
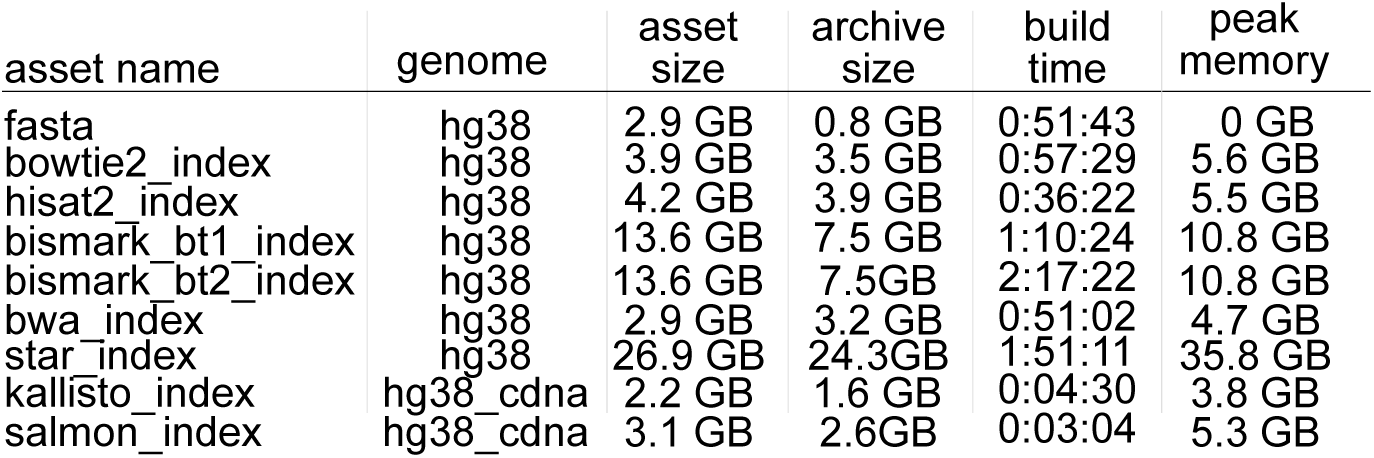
Assets available for build. Table listing assets that can currently be built with refgenie build, along with statistics for size, build time, and memory high water mark. Assets were built for the human genome using a single core. Times and memory are representative values from a single run. These assets are produced by various tools^8,9,14–17^ and are available to be built for any arbitrary genome input.

- `refgenie listr` lists available remote genomes and assets
- `refgenie pull` downloads a remote asset

With these commands, refgenie makes downloading a standard index for a user as simple as a few lines of code in a shell. For example, a new user can initialize refgenie and then download the bowtie2 indexes for the hg38 reference with these lines of code:

~~~
pip install --user refgenie
refgenie init -c conf.yaml
refgenie pull -c conf.yaml -g hg38 -a ASSET
~~~

Where `ASSET` is a unique key defining the asset of interest (*e.g.*, `bowtie2 index`). Once the asset has been pulled (or built), the user can retrieve the path to it with `refgenie seek`:

~~~
refgenie seek -c config.yaml -g hg38 -a ASSET
~~~

This command returns the file path to the specified asset for the specified genome. This command is now portable, eliminating the need to hard-code paths, or pass them as arguments, in a pipeline or other software that requires access to genome assembly assets.

### Refgenie server

The `listr` and `pull` functions require that the CLI interact with a server. The CLI uses a configurable URL to retrieve a remote archived tarball. After retrieving the tarball, the CLI will unpack it into the appropriate folder location and update the configuration file to provide access to its path via `refgenie seek`.

To support this remote function, we have developed a containerized, portable, open-source companion application called `refgenieserver`. Many users of refgenie will not have to be aware of the server application; however, interested users can use refgenie server to host their own genome asset server. For example, a tool developer may wish to simplify use by hosting indexes for common reference assemblies.

Running the refgenie server is simple for users who are already familiar with refgenie. It reads the same genome configuration file format as the CLI (indeed, it uses the `refgenconf` package described earlier in the same way). In fact, refgenie server operates on the same genome config file and asset folders that that refgenie itself builds or downloads. The server software comes with an `archive` command that prepares a refgenie genome folder for serving. It compresses each asset into an individual tarball. This simple system makes it easy for users to run a server using their refgenie assets.

This server software leverages cutting-edge web technology to provide high-concurrency service with minimal compute resources (Fig. 4). We built refgenie server on top of the FastAPI python framework, which is a high performance web framework for building APIs. FastAPI automatically produces an API that complies with OpenAPI 3.0 standards, which will allow other tools to discover and automatically use the API. It also includes a self-documenting test interface so that users can see and test the available API endpoints. Refgenie leverages the Starlette development toolkit and the `uvicorn` server to make use of the lightning-fast Asynchronous Server Gateway Interface (ASGI) specification, which provides asynchronous access to refgenie server.

**Fig.4:**
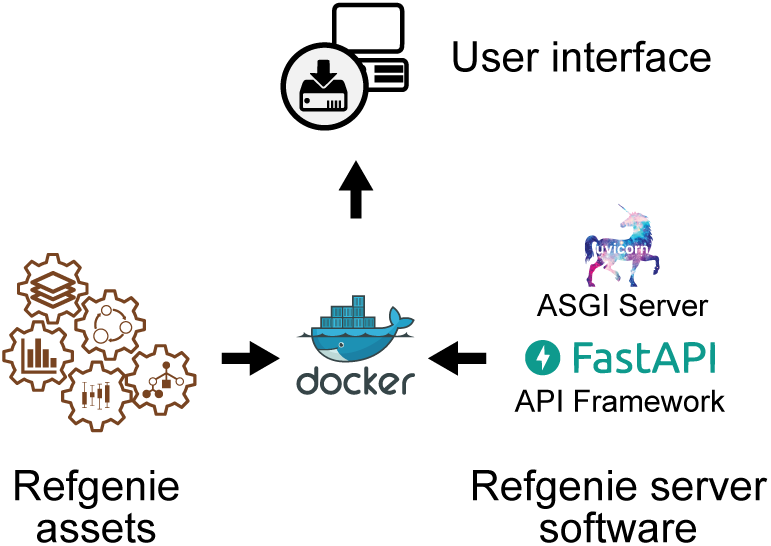
Server software stack. Archived refgenie assets are mounted into a Docker container, along with the refgenie server software, which is built using FastAPI and uvicorn. The container can then be accessed via the web and API user interfaces.

Refgenie server is containerized and available on dockerhub, so that an interested user could run a server with a single line of code:

~~~
docker run --rm -p 80:80 \
 -v genomes_folder:/genomes rgimage \
 refgenie -c /genomes/config.yaml serve
~~~

By mounting a refgenie ‘genomes’ folder into this container, users get a fully functioning web interface and RESTful API.

### The Refgenomes database

We designed the server software so that anyone could easily run a custom server instance. We have also deployed our own instance of `refgenieserver` at refgenomes.databio.org, where we host pre-built genome assets. Like any instance of `refgenieserver`, our refgenomes database provides both a web interface and a RESTful API to access individual assets we have made available. Users may explore and download archived indexes from the web interface or develop tools that programmatically query the API.

The web interface provides a graphical listing of available genomes and assets, allowing users to browse the site and download individual assets manually. In addition, refgenieserver provides API endpoints to serve lists of available genomes and assets, as well as metadata for the individual assets, including checksums for file integrity, file sizes, and archive content information. Furthermore, the server provides endpoints to download each asset individually. Endpoints include the following: `/genomes` retrieves a list of available genomes; `/assets` retrieves a list of all available assets; `/ { genome }/assets/` retrieves a list of assets for a given genome; and `/ {genome} /assets/ {asset} /archive` retrieves the tarball for the specified asset. Complete documentation is available at refgenomes.databio.org. Because it provides a standard OpenAPI-compliant RESTful API, our server will be useful not just for our refgenie CLI, but for other tools that would benefit from automated access to reference assembly assets and indexes.

Our `refgenieserver` instance runs within DC/OS as a containerized application managed by Marathon. Marathon deploys each application stack separately, monitors individual container health, and connects them to remote NFS storage and HTTP load balancers as appropriate. Marathon also has the ability to auto-scale cluster deployments. The refgenie application makes genome assets available through a web application connected directly to a remote filesystem, with no additional database or infrastructure requirements. Integration and deployment of frequently updated components is automated using GitHub, Travis-CI, Docker Hub, and a custom deployment technique made simple in DC/OS. Changes committed in code are generally deployed to development or production services within 1-3 minutes.

## Conclusions

Reference genomes, indexes, annotations, and other genome assets are integral to sequencing analysis projects, and these genome-associated data resources are growing rapidly^11^. Refgenie provides a full-service management system that includes a convenient method for downloading, building, sharing, and using these resources. Refgenieserver is among a growing number of API-oriented projects in the life sciences^5,18,19^. Refgenie will simplify management of reference assembly assets for users and groups, facilitating data sharing and software interoperability^20^.

## Notes

http://refgenie.databio.org

## References

1. Harrow, J. et al. GENCODE: The reference human genome annotation for the ENCODE project. Genome Research 22, 1760–1774 (2012).

2. Pruitt, K. D., Tatusova, T., Brown, G. R. & Maglott, D. R. NCBI reference sequences (RefSeq): Current status, new features and genome annotation policy. Nucleic Acids Research 40, D130–D135 (2011).

3. Church, D. M. et al. Modernizing reference genome assemblies. PLoS Biology 9, e1001091 (2011).

4. Kitts, P. A. et al. Assembly: A resource for assembled genomes at NCBI. Nucleic Acids Research 44, D73–D80 (2015).

5. Ruffier, M. et al. Ensembl core software resources: Storage and programmatic access for DNA sequence and genome annotation. Database 2017, (2017).

6. Sadakane, K. & Shibuya, T. Indexing huge genome sequences for solving various problems. Genome Informatics 12, 175–183 (2001).

7. Hon, W.-K., Sadakane, K. & Sung, W.-K. Breaking a time-and-space barrier in constructing full-text indices. SIAM Journal on Computing 38, 2162–2178 (2009).

8. Li, H. & Durbin, R. Fast and accurate short read alignment with burrows-wheeler transform. Bioinformatics 25, 1754–60 (2009).

9. Langmead, B. & Salzberg, S. L. Fast gapped-read alignment with bowtie 2. Nat. Methods 9, 357–359 (2012).

10. Illumina. IGenomes. Ready-to-use reference sequences and annotations. support.illumina.com (2019).

11. Richa Agarwala et al. Database resources of the national center for biotechnology information. Nucleic Acids Research 46, D8–D13 (2018).

12. Zerbino, D. R., Wilder, S. P., Johnson, N., Juettemann, T. & Flicek, P. R. The Ensembl Regulatory Build. Genome Biology 16, (2015).

13. Sheffield, N. C. & Bock, C. LOLA: Enrichment analysis for genomic region sets and regulatory elements in R and bioconductor. Bioinformatics 32, 587–589 (2016).

14. Krueger, F. & Andrews, S. R. Bismark: A flexible aligner and methylation caller for bisulfite-seq applications. Bioinformatics 27, 1571–1572 (2011).

15. Bray, N. L., Pimentel, H., Melsted, P. & Pachter, L. Near-optimal probabilistic RNA-seq quantification. Nature Biotechnology 34, 525–527 (2016).

16. Kim, D., Langmead, B. & Salzberg, S. L. HISAT: A fast spliced aligner with low memory requirements. Nature Methods 12, 357–360 (2015).

17. Dobin, A. et al. STAR: Ultrafast universal RNA-seq aligner. Bioinformatics 29, 15–21 (2012).

18. Yates, A. et al. The ensembl REST API: Ensembl data for any language. Bioinformatics 31, 143–145 (2014).

19. Tarkowska, A. et al. Eleven quick tips to build a usable REST API for life sciences. PLOS Computational Biology 14, e1006542 (2018).

20. Wilkinson, M. D. et al. The FAIR guiding principles for scientific data management and stewardship. Sci. Data 3, 160018 (2016).

